# Glucoregulatory Reprogramming in Male Mice Offspring Induced by Maternal Transfer of Indoor Flame Retardant Endocrine Disruptors

**DOI:** 10.1101/2022.10.02.510485

**Authors:** Elena V. Kozlova, Bhuvaneswari D. Chinthirla, Anthony E. Bishay, Pedro A. Pérez, Maximillian E. Denys, Julia M. Krum, Nicholas V. DiPatrizio, Margarita C. Currás-Collazo

## Abstract

Polybrominated diphenyl ethers (PBDEs) are commercially used as indoor flame retardants that penetrate biota and bioaccumulate in human tissues, including breast milk. PBDEs have been associated with endocrine disruption, diabetes and metabolic syndrome (MetS) in humans and animals. However, their sex-specific diabetogenic effects are not completely understood. Our past works show diabetogenic effects of the commercial penta-mixture of PBDEs, DE-71, in perinatally exposed C57Bl/6 female mice. As a comparison, in the current study, the effects of DE-71 on glucose homeostasis in male offspring were examined. C57BL/6 dams were exposed to DE-71 at 0.1 mg/kg/d (L-DE-71), 0.4 mg/kg/d (H-DE-71) or received corn oil vehicle (VEH/CON) for a total of 10 wks, including gestation and lactation. Male offspring were examined in adulthood and DE-71 exposure produced hypoglycemia upon extended fasting. *In vivo* glucose challenge testing showed marked intolerance (H-DE-71) and incomplete clearance (L- and H-DE-71). Moreover, L-DE-71-exposed mice showed altered glucose responses to insulin, especially incomplete glucose clearance and/or utilization. In addition, L-DE-71 produced elevated levels of plasma glucagon and the incretin GLP-1 but no changes were detected on insulin. These alterations, which represent relevant criteria used clinically to diagnose diabetes, were accompanied with reduced hepatic glutamate dehydrogenase enzymatic activity, elevated adrenal epinephrine and decreased thermogenic brown adipose tissue mass, which may indicate several organ system targets of PBDEs. Liver levels of several endocannabinoid species were not altered by perinatal exposure to DE-71 in males. Our findings demonstrate that chronic low exposure to PBDEs in mothers can reprogram glucose homeostasis and glucoregulatory hormones in male offspring. Previous findings using female offspring showed altered glucose homeostasis that aligned with a contrasting diabetogenic phenotype. We summarize the results of the current work generated in males in light of previous findings on females. Taken together, these findings, combined with our prior results, offer a comprehensive account of sex-dependent effects of maternally transferred environmentally relevant PBDEs on glucose homeostasis and glucoregulatory endocrine dysregulation.

## Introduction

The recent epidemic rise in metabolic disease, obesity and type 2 diabetes (T2D), liver lipid disorder and metabolic syndrome (MetS) cannot solely be attributed to genetic background and lifestyle changes. Considerable evidence points to a potential contribution from endocrine-(EDCs) and metabolism-disrupting chemicals (MDCs) to the rapid increase in the incidence of these metabolic diseases (1). One class of EDCs/MDCs are polybrominated diphenyl ethers (PBDEs), anthropogenic persistent organic pollutants (POPs), that have been widely used as flame retardants in commercial products such as furniture, carpets, automobiles, building materials and electronics (2) since the 1970’s. PBDEs are lipophilic additives which are not chemically bound and can be readily released into the environment and bioaccumulate in biota. PBDEs form 209 theoretical congeners depending on the number of bromine substitutions on the biphenyl backbone, with tetra-, penta- and hexa-substituted PBDEs being most commonly found in humans. Research in experimental animals and epidemiological studies has revealed that PBDEs impart toxicological actions on reproduction (3), (4) neurodevelopment (5) and thyroid homeostasis (6). While these findings have led to a ban on the production and usage of PBDEs by the European Union in 2004 and to the voluntary phase out of in the US starting in 2005, biomonitoring data indicates PBDE are still found in significant amounts in the placenta (7), fetal blood (8) and breast milk (9). Moreover, deca-PBDEs are still used in television casing and other products, and these can be debrominated to more harmful penta- and other derivatives (10). Therefore, modeling studies show that PBDEs levels are predicted to be present in the environment until 2050 due to inadvertent recycling and e-waste (11).

The T2D epidemic has become a serious global health issue, affecting 415 million people world-wide (12) and poses a significant economic burden on individuals and society (13). A large body of evidence from humans and animal research implicates brominated flame retardants such as PBDEs in the pathogenesis of T2D and MetS (14), (15), (16), (17), (18), (19), (20), (21), (22). A commonality in all these studies is that diabetes and/or MetS are positively associated with body burdens of the PBDE congeners BDE-28, −47, and −153, all found in humans and biota. The PBDE congeners that have been associated with T2D are found in DE-71, a commercial mixture of PBDEs. Our past work shows that DE-71 congeners at the doses used here can be transferred perinatally via the dam and accumulate in offspring liver (23) and brain (24) at environmentally relevant levels. DE-71 may lead to metabolic reprogramming in offspring especially if exposure is administered during a period of high biological plasticity such as gestation and lactation. Indeed, we have shown that female offspring exposed to 0.1 mg/kg/d DE-71 perinatally, alters clinically relevant biomarkers of T2D - fasting blood glucose, glucose tolerance, insulin sensitivity, plasma levels of glucoregulatory hormones and wells as liver endocannabinoid tone, an emerging biomarker of energy balance. Further, we have also found that PBDEs can affect hypothalamic peptidergic circuits and, namely, can disrupt circuits that control food intake and energy metabolism in a sex dependent manner (Kozlova et al., 2022, *In press*).

The interplay between the key endocrine regulators of glucose homeostasis, glucagon, insulin, and glucagon-like-peptide (GLP-1) is of critical importance in understanding the diabetogenic phenotype of individuals with T2D. The incretin GLP-1 is secreted by enteroendocrine L-cells and acts on pancreatic *β*-cells to promote postprandial insulin secretion. In the T2D, this insulinotropic effect is either impaired or absent (25). GlP-1 receptors are also found in the hypothalamus where GLP-1 promotes satiety and in pancreatic α-cells where it suppresses glucagon secretion. Glucagon, released from pancreatic α-cells in the fasting state, acts reciprocally to insulin to promote normoglycemia. However, in T2D there is a lack of inhibitory tone of insulin on glucagon secretion leading to elevated glucagon levels (26). We have previously shown the differential effect of PBDEs on insulin and glucagon in female offspring and GLP-1 in dams exposed in adulthood (23). While few studies have examined sex-differences in T2D patients (27) and less is known about sex differences in hormonal pathophysiology of T2D, several hormones regulating glucose control have been found to vary by sex and body type (28). EDCs, including PBDEs, have been shown to alter insulin levels (29), (30) and increase the risk of gestational diabetes (31). However, reports of the sex-related differences on the interplay between the major glucoregulatory endocrine hormones as a result of environmental toxicants, like PBDEs, is unclear. Moreover, EDCs involvement in T2D has not been studied using an integrative physiological approach that includes multiple organ systems such as liver and sympathetically innervated brown adipose tissue.

Having observed sexually dimorphic risk and susceptibility of perinatally exposed females to MetS, the purpose of this study was to comprehensively examine the effect on glucose homeostasis in male offspring. Using a mouse model of chronic, low-dose maternal transfer of environmentally relevant PBDE congeners, we tested the hypothesis that exposure to DE-71 produces a diabetogenic phenotype in male offspring. Results indicate that exposure to DE-71, alters clinically relevant diabetic biomarkers in males, namely, fasting blood glucose, glucose tolerance, insulin sensitivity, plasma levels of glucoregulatory hormones, however, clear sex differences emerge. Taken together, the current study and our past work elucidates a comprehensive profile of the clinically relevant and persistent metabolic reprogramming induced by PBDEs and raises concern for the progeny of directly exposed mothers.

## Methods

### Animals

C57Bl/6N mice were obtained from Charles River (Raleigh, NC) or Taconic Biosciences (Germantown, NY). Mice were group housed 2-4 per cage and maintained in a non-specific pathogen free vivarium on a 12 h light/dark cycle at an ambient temperature (20.6-23.9°C) and relative humidity environment (20-70%). Mice were provided rodent chow (Laboratory Rodent Diet 5001; LabDiet, USA) and municipal tap water *ad libitum* in glass water bottles. Procedures on the care and treatment of animals were performed in compliance with the National Institutes of Health *Guide for the Care and Use of Laboratory Animals* and approved by the University of California, Riverside Institutional Animal Care and Use Committee (AUP#20170026 and 20200018).

### Dosing Solutions

Dosing solutions were prepared as described previously (23), (24). In brief, technical pentabromodiphenyl ether (DE-71; Lot no. 1550OI18A; CAS 32534-81-9), was obtained from Great Lakes Chemical Corporation (West Lafayette, IN). DE-71 dosing solutions were prepared to yield two ultra-low doses: 0.1 (L-DE-71) and 0.4 (H-DE-71) mg/kg bw per day 2 mL of stock solution/kg body weight. Vehicle control solution (VEH/CON) contained corn oil without the addition of DE-71.

### DE-71 Exposure

Perinatal PBDE exposure via the dam was accomplished as described previously (23). In brief, female mice (PND 30-60) were introduced to cornflakes daily for 1 week. Dams were randomly assigned to one of three exposure groups: corn oil vehicle control (VEH/CON), 0.1 (L-DE-71) or 0.4 mg/kg (H-DE-71) bw per day DE-71 (**Fig. 1**). A 10-week dosing regimen, chosen to model human-relevant chronic, low-level exposure (32), (33), (34), included ~4 weeks of pre-conception, plus gestation (3 weeks) and lactation (3 weeks). Offspring were weaned after the lactation period at PND 21 and housed in same-sex groups. Dams were fed oral treats, (Kellogg’s Corn Flakes) infused with dosing solution (2 uL/g bw) daily, except on PND 0 and 1. Each dam was exposed to DE-71 daily for an average of 70-80 d and offspring were perinatally exposed for 39 d via maternal blood and milk. Consumption was visually confirmed, and offspring were never allowed to ingest cornflakes. DE-71 exposure does not significantly affect maternal food intake, gestational weight gain, litter size or secondary sex ratio (23). Male offspring were used *in vivo* and *ex vivo* for analysis of physiological, metabolic, and endocrine parameters. The DE-71 exposure and testing paradigm is shown in **Figure 1**.

**Figure 1.**
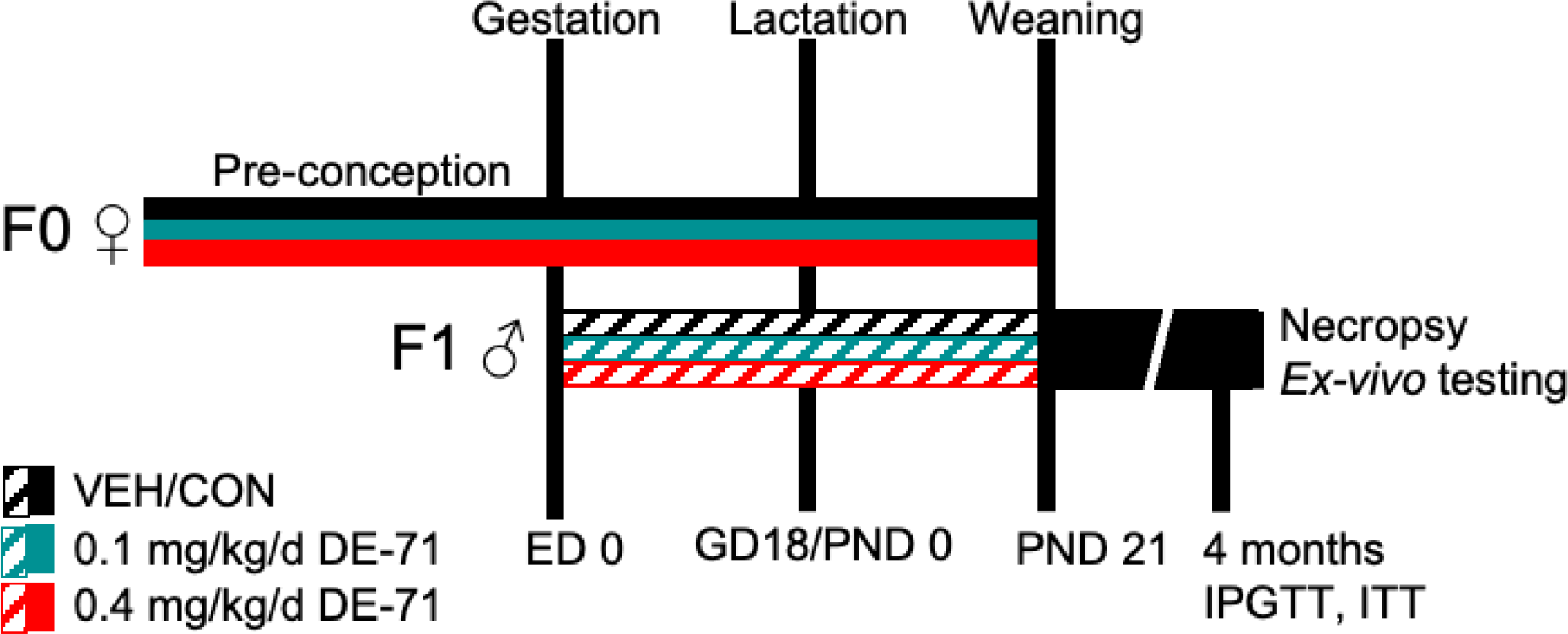
Diagram depicting the dosing and testing paradigm used for perinatal exposure to DE-71. Direct exposure of dams to DE-71 (F0♀, solid shading), began ~4 weeks pre-conception and continued daily through ~3 weeks of gestation and 3 weeks of lactation until pup weaning at PND 21. Indirect exposure to DE-71 in male offspring (F1♂, hatched shading) occurred during the perinatal (ED 0-GD 18) and postnatal periods (PND0-PND21) via lactation. Offspring received 39 consecutive days of DE-71 exposure. Metabolic endpoints (fasting glycemia, GTT, ITT) were examined in offspring at ~4 months of age. At necropsy, body and organ weights were recorded and blood and organ tissues collected for further analysis: plasma hormones, UPLC/MS/MS detection of endocannabinoids, adrenal epinephrine and liver GDH enzymatic activity. GD, gestational day; PND, postnatal day; ED, embryonic day; GDH, glutamate dehydrogenase.

### Necropsy

During sacrifice, under terminal isoflurane anesthesia, cardiac blood (0.3-1 mL) was collected and centrifuged at 16,000 x g for 20 min at 4 °C. After the addition of A cocktail of protease inhibitors and EDTA, the plasma samples were stored at −80 °C until further use. The following organs were excised and weighed: liver, pancreas, spleen, adrenal glands and interscapular BAT. Plasma, liver and adrenal samples were snap-frozen over dry ice and stored at −80 °C for later analysis of plasma hormones, adrenal epinephrine, liver endocannabinoids and enzymatic activity.

### Glucose Tolerance test (GTT) and Insulin Tolerance Test (ITT)

Mice were fasted overnight (ON) for 11h and then injected with glucose (2.0 g/kg bw, i.p.). Following the collection of tail blood (1 uL), glucose was sampled at time 0 (FBG) 15, 30, 60, and 120 min post-glucose challenge. A calibrated glucometer (OneTouch Ultra 2, LifeScan Inc.) was used to measure plasma glucose concentrations. Seven days following GTT, an insulin tolerance test (ITT) was performed with Humulin R bolus (0.25 U/kg bw, i.p.) on mice fasted ON for 9h. Tail blood was collected, and glucose was sampled in the same manner as IPGTT (t=0 (FBG), 15, 30, 45, 60, 90 and 120). For area (AUC) calculations, the area under or above the glycemia curve (inverse AUC) from 0-120 min post injection was used. The blood glucose reduction rate after insulin administration, K_ITT_, was calculated using the formula 0.693 × t_1/2_^−1^ to determine *in vivo* insulin sensitivity from 0-30 min post insulin injection.

### Immunoassays

Plasma collected via cardiac puncture (*ad libitum* fed state) was analyzed using commercially available kits according to manufacturer’s instructions. Plasma insulin was measured using commercial ELISA kits (ALPCO, Salem, NH, Cat.# 80-INSMS-E01 and Mercodia, Uppsala, Sweden Cat.# 10-1249-01 and 10-1247-01) as previously described(23). The Mercodia assays had a sensitivity of 0.15 mU/L or 1 mU/L in a standard range of 0.15-20 mU/L or 3-200 mU/L. The ALPCO insulin ELISA had a sensitivity of 0.019 ng/mL in a standard range of 0.025-1.25 ng/mL and inter- and intra-assay CV of 5.7% and 4.5%, respectively. The active glucagon-like peptide-1 (7-36) Amide (GLP-1) assay (Cat.# 80-GLP1A-CH01, ALPCO) had an analytical sensitivity of 0.15 pM in a standard range of 0.45-152 pM and inter- and intra-assay CV of 11.6 and 9.5%, respectively. Glucagon was measured by chemiluminescence ELISA (ALPCO, Cat.# 48-GLUHU-E01). This assay had a sensitivity of 41 pg/mL in a dynamic range of 41-10,000 pg/mL and inter- and intra-assay CV of 9.8% and 7.6%, respectively.

### Ultra-performance liquid chromatography-tandem mass spectrometry

Hepatic lipids were extracted using a modified Folch method as described previously (23). In brief, samples of flash-frozen liver tissue were weighed (10-20 mg) and homogenized in 1 mL of methanol solution containing 1 pmol d_4_-arachidonoylethanolamide, 10 pmol d_4_-oleoyloethanolamide, and 500 pmol d_5_-2-arachidonoyl-*sn*-glycero as internal standards. Following homogenization, 2 mL of chloroform and 1 mL of water were added, and centrifuged at 2000 x g for 15 min at 4 °C. The organic phase was extracted with chloroform. The pooled lower phases were dried under N_2_ gas followed by resuspension in 0.1 mL methanol:chloroform (9:1). Analysis of EC was performed via ultra-performance liquid chromatography coupled to tandem mass spectrometry (UPLC/MS/MS) as previously described (35).

### Glutamate dehydrogenase (GDH) activity

GDH activity in crude liver homogenates was assayed using the tetrazolium salt method with modification for multiwell plates as described previously (23). In brief, 5% liver homogenates (40 uL in 0.25M sucrose) were added to a mixture of 2 μmol/l of iodonitrotetrazolium chloride, 0.1 μmol/l of NAD, 50 μmol/l of sodium glutamate, 100 μmol/l of phosphate buffer (pH 7.4) and distilled water. Samples were run in duplicate. The reaction was allowed to proceed at 37 °C for 30 min. The resultant formazan product was measured as optical density at 545 nm and then converted to concentration using a iodonitrotetrazolium formazan (TCI) standard curve fitted with a linear regression model (36). A bicinchoninic acid assay (Cat.# 23227, ThermoFisher Scientific, Waltham, MA, USA) was used to measure protein content in order to normalize product values. GDH activity was expressed as μmol formazan formed per ug protein/h.

### Epinephrine Assay

Epinephrine content in adrenal glands was measured using a modification of the trihydroxyindole method as described previously (23). Briefly, 5 mg of adrenal tissue homogenates (in 200 μL of 0.05 N perchloric acid) were centrifuged at 15,000 x g at 0 °C for 15 min. 10% acetic acid (pH 2) was added to the sample supernatant (30 uL), followed by the addition of 60 μL of 0.25% K_2_Fe(CN)_6._. The mixture was incubated at 0 °C for 20 min and the oxidation reaction was stopped by the addition of 60 μL of a 9 N NaOH solution containing 4 mg/ml ascorbic acid (alkaline ascorbate). Epinephrine concentration (μg/g adrenal wet weight) was converted from the mean fluorescence intensity units of each sample using calibration standards and polynomial curve fitting.

### Statistical Analysis

Statistical analyses were performed using GraphPad Prism v.9.4.1. A one-way analysis of variance (ANOVA) was used to test the main effect of one factor. When normality assumption failed, determined using a Shapiro-Wilk test, a Kruskal-Wallis H ANOVA was used. A Brown-Forsythe ANOVA was used if the group variances were significantly different. Data for fasting glycemia were analyzed by two-way ANOVA for main effects of exposure and fasting duration. ITT and GTT experiments were analyzed by repeated measures two-way or mixed model ANOVA to determine main effects of exposure and time after challenge. ANOVA was followed by *post hoc* testing for multiple group comparisons. Differences were deemed significant at *p*<0.05. Data are expressed as mean±s.e.m.

## Results

### Chronic low dose DE-71 exposure has minimal effects on body weight, select organ weights and organ-to-body-weight-ratios

Table 1 shows effects of DE-71 exposure on body weights, select organ weights and organ-to-body-weight-ratios. Necropsy body weights of male offspring were not different across groups. Absolute liver weight was 9% lower in L-DE-71 relative to VEH/CON (One-way ANOVA: *Exposure effect* F_(2,58)_=9.34, *p*<0.001, Tukey’s *post-hoc* VEH/CON vs L-DE-71, *p*=0.03, L-DE-71 vs H-DE-71 *p*=0.0003). Relative liver weight was 8% greater in H-DE-71 relative to L-DE-71 (One-way ANOVA: *Exposure effect* F_(2,58)_=2.824, *p*=0.0675, Tukey’s *post-hoc* L-DE-71 vs H-DE-71 *p*=0.054). The absolute and relative weights of pancreas and spleen were similar across groups. In addition, body weights were not different across groups and, therefore, the diabetogenic phenotype of DE-71-exposed male mice is not due to obesity. This is consistent with our previous findings of no difference in lean or fat mass in DE-71-exposed males (Kozlova et al., 2022, *In press*)

**Table 1.** Body Weight, Select Organ Weights and Organ-Weight-to-Body-Weight Ratios in DE-71 Exposed Male Mice.

|  | VEH/CON | L-DE-71 | H-DE-71 |
| --- | --- | --- | --- |
| n | 17 | 28 | 18 |
| Necropsy body weight | 25.7 ± 0.50 | 24.8 ± 0.71 | 25.7 ± 0.49 |
| Liver |  |  |  |
| Absolute | 1.16 ± 0.03 | 1.06 ± 0.02* | 1.21 ± 0.03<sup>^^^</sup> |
| Relative | 44.8 ± 1.70 | 43.2 ± 1.05 | 47.1 ± 1.05<sup>^</sup> |
| Pancreas |  |  |  |
| Absolute | 0.17 ± 0.007 | 0.17 ± 0.007 | 0.19 ± 0.008 |
| Relative | 6.37 ± 0.26 | 6.94 ± 0.21 | 7.27 ± 0.30 |
| Spleen |  |  |  |
| Absolute | 0.07 ± 0.002 | 0.07 ± 0.002 | 0.07 ± 0.003 |
| Relative | 2.64 ± 0.10 | 2.85 ± 0.10 | 2.80 ± 0.09 |

Values for body and organ weights (absolute weights) are expressed in grams; organ-weight-to-body-weight ratios (relative weights) are given as mg organ weight/g body weight. Values are reported as mean ± s.e.m.
<sup>^</sup> Indicates significantly different relative to L-DE-71 (<sup>^</sup> $p$ <0.05; <sup>^^^</sup> $p$ <0.001)

### DE-71 produces fasting hypoglycemia

We examined FBG after 9 and 11 h fasting using glycemia values from basal time points obtained in ITT and GTT experiments, respectively. Figure 2 shows that exposure to H-DE-71 significantly lowered (25%) FBG after a 11 h fast relative to VEH/CON (Two-way ANOVA: *Exposure effect* F_(2,78)_= 6.65, *p*<0.01; *Time effect* F_(1,78)_= 24.6, *p*<0.0001; *Exposure* x *Time* F_(6,78)_= 0.76; Sidak’s *post hoc* test, 11 h: VEH/CON vs L-DE-71, *p*<0.01; 9 h vs 11 h, L-DE-71 *p*<0.01, H-DE-71, *p*<0.05 (**Fig. 2**). In comparison, this was not observed after a shorter 9 h fast. These results suggest that perinatal exposure to DE-71 alters fasting glycemia, which may be due to dysregulated endocrine hormones involved in glucose metabolism (37).

**Figure 2.**
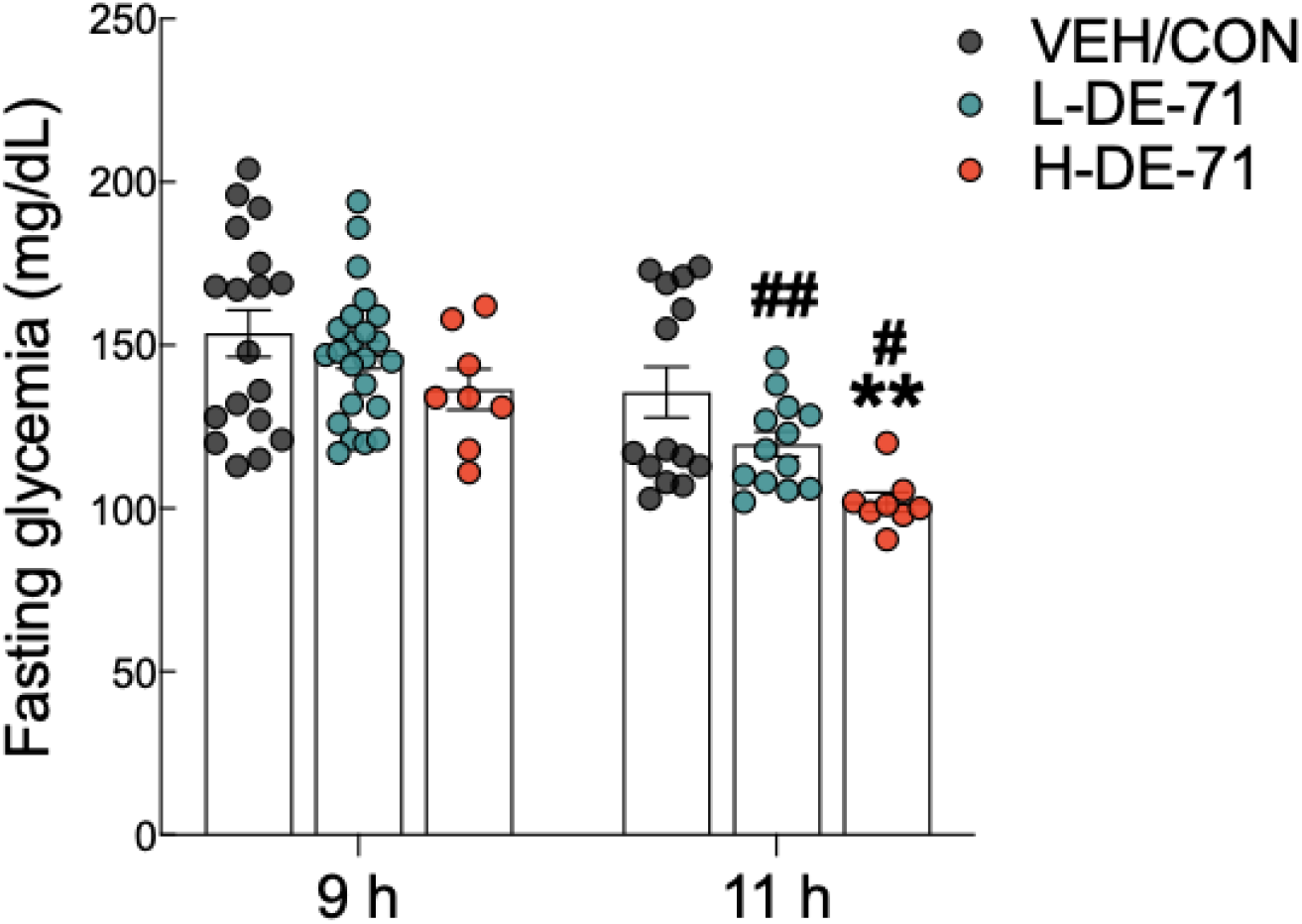
Fasting blood glucose (FBG) in male offspring exposed perinatally to DE-71. FBG was measured after a 9 and 11h fast in male offspring. *indicates significantly different from VEH/CON (\*\**p*<0.01). ^#^indicates significant difference between 9 and 11 h for corresponding exposure group (#*p*<0.05; ##*p*<0.01). Bars and error bars represent values expressed as mean±s.e.m. *n*, 8-23/group.

### DE-71 exposure impairs glucose tolerance

To investigate the effects of DE-71 on glucose tolerance, glycemia was measured during GTT over the 120 min post-injection time course (**Fig. 3A**). Blood glucose levels rose rapidly and peaked between 15-30 min of glucose challenge in all groups. Relative to VEH/CON, peak glycemia was exaggerated in H-DE-71 males relative to VEH/CON (RM Two-way ANOVA: *Time effect* F_(2.9,96.1)_=330.2, *p*<0.0001; *Exposure effect* F_(2,33)_=4.767, *p*<0.05; *Time* x *Exposure* F_(8,132)_=3.112, *p*<0.01) (**Fig. 3A**). Mean glycemia values in H-DE-71 were markedly elevated as compared to VEH/CON at *t*=30 (*p*<0.05) and 60 min post injection (*p*<0.05) and to L-DE-71 at *t*=30 (*p*<0.01) and 60 min post injection (*p*<0.01).

**Figure 3.**
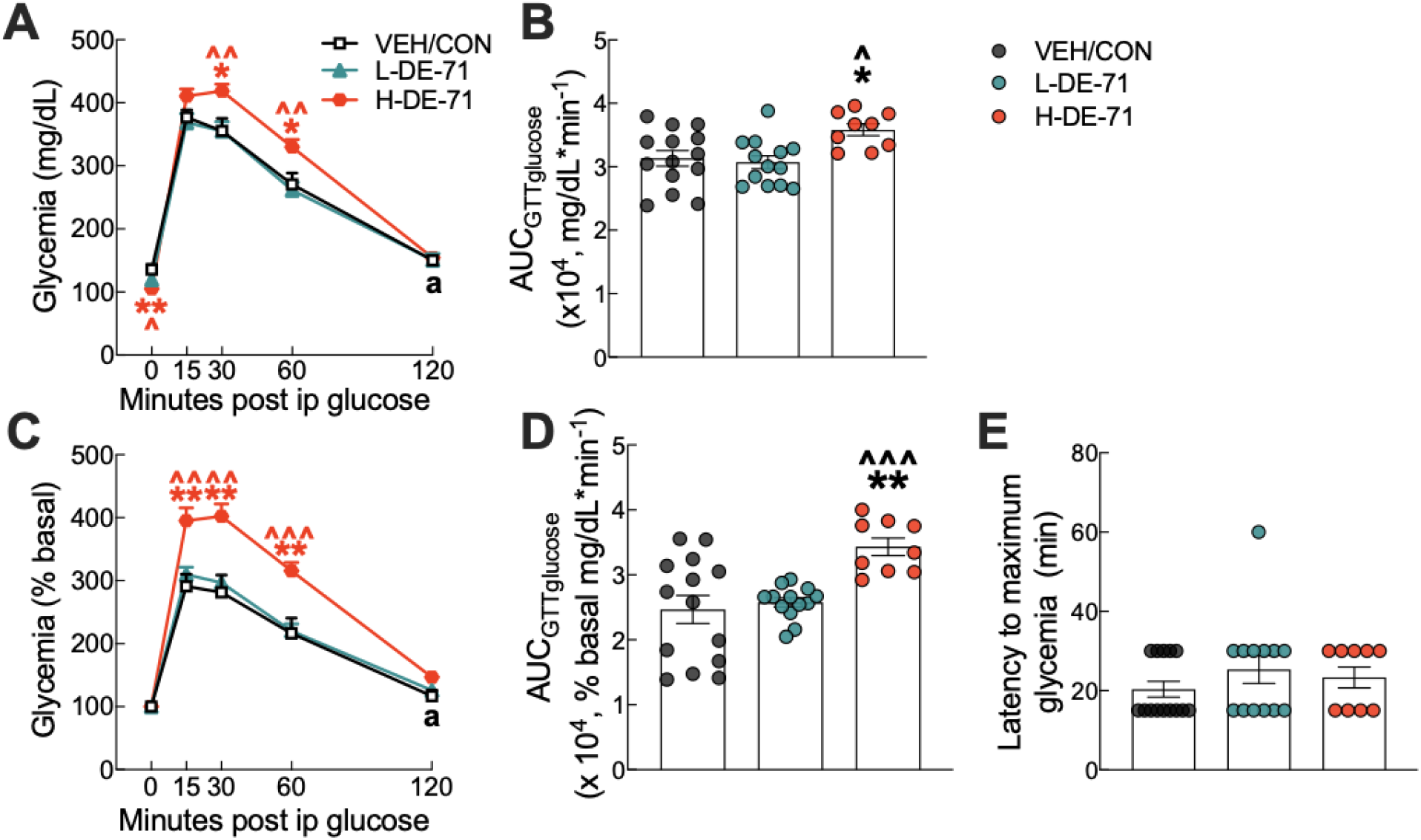
Glucose tolerance profile in male mice offspring receiving perinatal exposure to DE-71. Mice were fasted for 11 h ON and tail blood was sampled for glucose before (t=0 min) and after (t=15, 30, 60 and 120 min) i.p. injection of 2.0 g/kg glucose. (**A**) Absolute blood glucose concentrations taken during IPGTT. (**B**) Mean values for the integrated area under the IPGTT glucose curve using absolute values (AUC_GTTglucose_). (**C**) Blood glucose values taken during IPGTT are plotted vs time as a percent of basal glucose. (**D**) Mean values for the integrated area under the IPGTT glucose curve using percent baseline values (AUC_GTTglucose_). (**E**) Latency to maximum glycemia corresponding to percent basal. Asterisk of specific color indicates significant difference between corresponding group and VEH/CON (\**p*<0.05, \*\**p*<0.01). Carets of specific color indicate significant difference between corresponding group and L-DE-71 (^*p*<0.05, ^^*p*<0.01, ^^^*p*<0.001). The black symbol “a” in panels A and D indicates the time points at which glycemia is not different from basal in the VEH/CON group. Glycemia at all other time points differs from basal. Bars and error bars represent values expressed as mean±s.e.m. *n*, 9-14/group.

Since FBG after an 11 h fast (*t*=0) was elevated in H-DE-71 relative to VEH/CON (Fig. 2), glycemia values are also expressed as percent baseline (**Fig. 3C)**. After normalizing to baseline (t=0), peak glycemia was exaggerated in H-DE-71 males relative to VEH/CON (RM Two-way ANOVA: *Time effect* F_(2.26, 74.7)_=278.6, *p*<0.0001; *Exposure effect* F_(2,33)_=9.39 *p*<0.001; *Time* x *Exposure* F_(8,132)_=5.94, *p*<0.0001). Mean glycemia values for H-DE-71 were significantly different from VEH/CON at *t*=15 (*p*<0.01), 30 (*p*<0.01) and 60 min post injection (*p*<0.01) and to L-DE-71 at *t*=15 (*p*<0.01), *t*=30 (*p*<0.01) and 60 min post injection (*p*<0.001).

The differences in magnitude and duration of glycemia are integrated using the area under the glucose curve, AUC_GTTglucose_, which is abnormally large in H-DE-71 males (Absolute glycemia AUC, One-way ANOVA: *Exposure effect* F_(2,33)_=5.21, *p*<0.05, Tukey’s *post-hoc* VEH/CON vs H-DE-71, *p*<0.029, L-DE-71 vs H-DE-71, *p*<0.013; Percent basal glycemia AUC, Brown-Forsythe ANOVA: *Exposure effect* F_(2,22.2)_=10.1, *p*<0.001, Dunnett’s T3 *post-hoc* VEH/CON vs H-DE-71, *p*<0.004, L-DE-71 vs H-DE-71 *p*<0.0003 (**Fig. 3B, 3D**). The latency to maximum glycemia was not significantly different across groups (**Fig. 3E**). Of note, glycemia was still elevated in both exposure groups (L- and H-DE-71) but not VEH/CON at 120 min post-glucose injection (“a” in Fig. 3a, c). These comparisons are still significant when comparing % basal glycemia at 120 min (L-DE71 p<0.05, H-DE-71 p<0.001). These results suggest that developmental exposure to DE-71 causes glucose intolerance (H-DE-71) and incomplete clearance (L- and H-DE-71).

### L-DE-71 exposure produces an abnormal glycemic response to insulin

The glycemia response to exogenous insulin was examined during ITT experiments. The insulin tolerance curve shows mean glycemia values over the 120 min period following insulin injection (**Fig. 4A)**. Males exposed to L-DE-71 display more reduction in glycemia as compared to VEH/CON (RM Two-way ANOVA: *Time effect* F_(3.257,143.3)_=56.2, *p*<0.0001; *Exposure effect* F_(2,44)_=2.40, *ns*; *Time* x *Exposure* F_(12,264)_=1.90, *p*<0.05) (**Fig. 4A**). Mean glycemia values for L-DE-71 were significantly different from VEH/CON at *t*=120 min (*p*<0.05). When glycemia is expressed as a percent of baseline, L-DE-71 exposed mice displayed incomplete recovery or glucose clearance/utilization after insulin injection (RM Two-way ANOVA: *Time effect* F_(6,308)_=33.7, *p*<0.0001; *Exposure effect* F_(2,308)_=8.014, *p*<0.001; *Time* x *Exposure* F_(12,308)_=1.23, *ns*) (**Fig. 4D**). Mean glycemia values for L-DE-71 are significantly different from VEH/CON at *t*=90 (*p*<0.05) and 120 min (*p*<0.001). This is represented as a greater mean latency to reach the minimum insulin-induced hypoglycemia in L-DE-71 relative to VEH/CON (Kruskal-Wallis test: Exposure H(2)=9.07, *p*<0.01; Dunn’s *post hoc* VEH/CON vs L-DE-71, *p*<0.01) (**Fig. 4F**).

**Figure 4.**
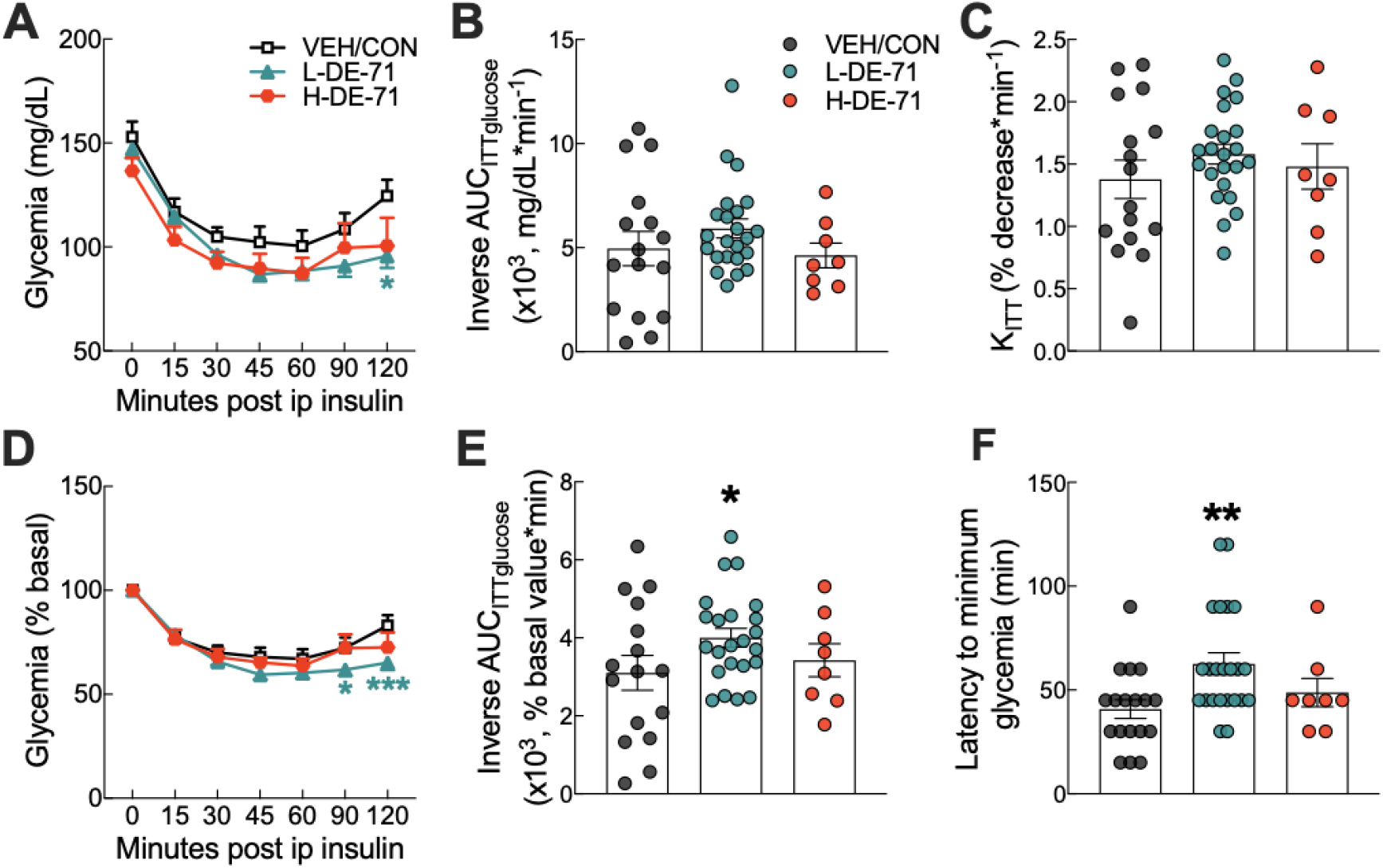
Abnormal glycemia reduction after insulin challenge in male mice offspring exposed perinatally to L-DE-71. (**A**) Absolute blood glucose concentrations were recorded before and at *t*=15, 30, 45, 60, 90 and 120 min post-injection with 0.25 U/kg insulin. (**B**) Tolerance to insulin challenge was analyzed by the inverse integrated area under the ITT glucose curve (AUC_ITTglucose_). (**C**) Rate constant for glucose reduction (K_ITT_) was calculated over the initial slope of ITT glucose response curve from 0-30 min post-injection. (**D**) Glucose values taken during ITT are plotted vs time as a percent of the individual baseline. (**E**) The inverse integrated area (AUC) under the percent basal glucose curve (AUC_ITTglucose_). (**F**) Latency to minimum blood glucose measured over the two-hour time course of ITT glucose response. *indicates significantly different from VEH/CON (\**p*<0.05, \*\**p*<0.01, \*\*\**p*<0.001). All values are expressed as mean±s.e.m. *n*, 8-23/group.

The inverse area under the glucose response curve (inverse AUC_ITTglucose_) was plotted using absolute (**Fig. 4B**) and percent baseline values (**Fig. 4E**), the latter showed a significant increase for L-DE-71 relative to VEH/CON, One-way ANOVA: *Exposure effect* F_(2,44)_=2.024, ns, Sidak’s *post-hoc* VEH/CON vs L-DE-71, *p*<0.05) (**Fig. 4E**). To measure the early effects of insulin sensitivity, we calculated K_ITTinsulin_ measured over the first 30 min post-injection (**Fig. 4C**). There were no differences between groups for this metric, suggesting no differences in early insulin clearance. Interestingly, male offspring of dams exposed to L-DE-71 during pregnancy are more sensitive to insulin characterized by a deeper trough on ITT graph. However, they also show a delayed recovery from insulin challenge, suggesting an altered metabolic phenotype.

### Endocrine-disrupting effects of DE-71 exposure on glucoregulatory hormones

We have previously shown disrupted plasma hormones involved in carbohydrate regulation in DE-71 exposed female offspring. In T2D, pancreatic beta cell dysfunction may lead to changes in insulin production(38). In addition, elevated fasting and postprandial plasma glucagon concentrations have also been shown and may contribute to symptomatic hyperglycemia, an effect that may be due to deficient GLP-1-mediated regulation(39). We used EIA to measure plasma insulin, GLP-1 and glucagon in blood collected at necropsy (*ad libitum* fed state) (**Fig. 5A**). Mean absolute concentrations ranged from 0.90 ± 0.12 to 1.12 ± 0.17 ug/L for insulin, 12.6 ± 4.98 to 41.9 ± 9.78 pg/mL for glucagon in F1 males. Contrary to our hypothesis, plasma insulin levels were not different across groups (**Fig. 5A**). However, L-DE-71 males showed elevated plasma GLP-1 and glucagon relative to VEH/CON (GLP-1, One-way ANOVA: *Exposure effect* F_(2,22)_=4.532, *p*=0.07; Dunnet’s *post-hoc* VEH/CON vs L-DE-71 *p*=0.02; Glucagon, One-way ANOVA: *Exposure effect* F_(2,15)_=2.210, *p*=0.14; Dunnett’s *post-hoc* VEH/CON vs L-DE-71 *p*=0.04) (**Fig. 5B,C**). In summary, L-DE-71 males showed the most endocrine susceptibility to PBDEs.

**Figure 5.**
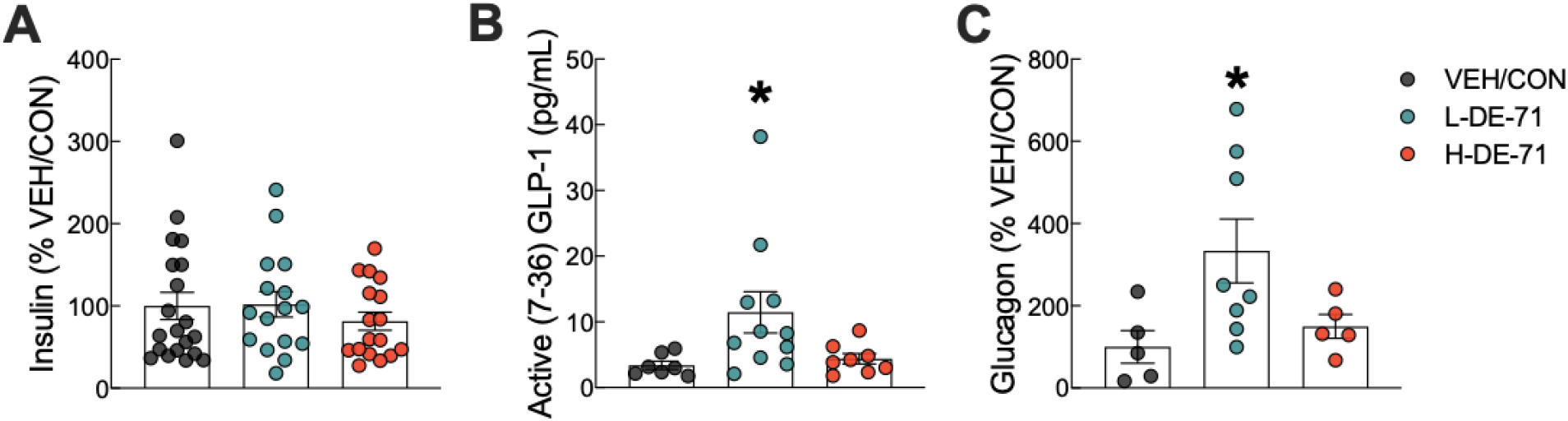
Endocrine-disrupting effects of DE-71 on glucoregulatory hormones in male mice. Blood collected at sacrifice was assayed for plasma levels of (**A**) insulin (**B**) GLP-1 and (**C**) glucagon using specific commercial enzyme-linked immunoassay kits. *indicates significantly different from corresponding VEH/CON (\**p*<0.05). Bars and error bars represent values expressed as mean±s.e.m. *n*=16-20/group for insulin, *n*=7-11/group for GLP-1, *n*=5-8/group for glucagon.

### Upregulated adrenal epinephrine content and reduced BAT after DE-71 exposure

Due to the important role of epinephrine in glucose and lipid homeostasis, we examined whether adrenal content of epinephrine was altered by developmental exposure to DE-71. Adrenal epinephrine levels were similar to those reported previously for male wildtype C57Bl/6 mice using a radioenzymatic assay and LC-MS measurement instead of fluoroscopy(40). Figure 6A shows that L- and H-DE-71 exposure significantly elevated adrenal epinephrine (One-way ANOVA: *Exposure effect* F_(2,4.213)_=3.312, *p*=0.0606; Tukey’s *post-hoc* VEH/CON vs L-DE-71 *p*=0.0002, VEH/CON vs H-DE-71 *p*=0.0022). Adrenal weights were not different across experimental groups (data not shown). Because diabetes and fasting glucose level is associated with brown adipose tissue (BAT) activity(41), we measured BAT mass(41). When normalized to body weight, mean interscapular BAT mass was dramatically decreased by 30.2% in L-DE-71 relative to VEH/CON (Brown-Forsythe ANOVA: *Exposure effect* F_(2,37.08)_=8.468, *p*=0.061; Dunnett’s *post-hoc* VEH/CON vs L-DE-71 *p*=0.049, L-DE-71 vs H-DE-71 *p*=0.004) (**Fig. 6B**).

**Figure 6.**
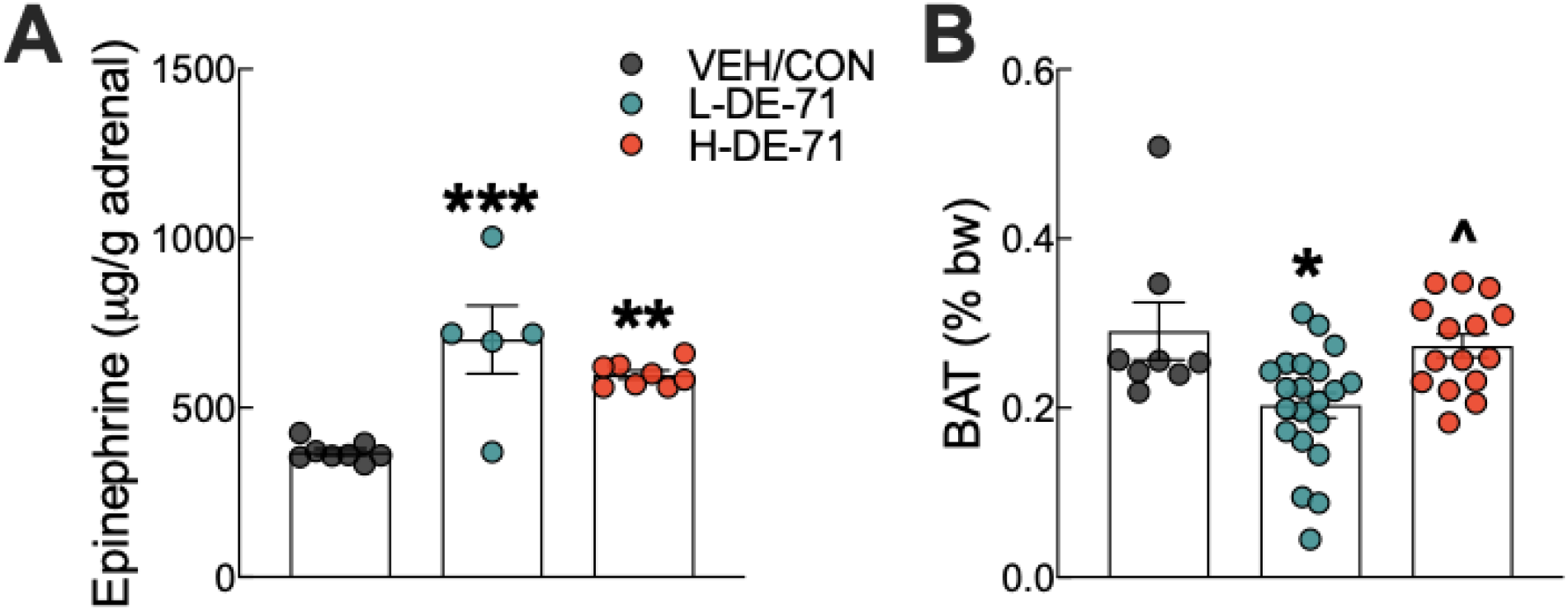
Effects of DE-71 on adrenal epinephrine content and thermogenic brown adipose tissue mass. **(A)** Epinephrine content measured in adrenal glands harvested at necropsy. (**B**) Interscapular brown adipose tissue (BAT) collected at necropsy was expressed as a percent of body weight. *indicates significantly different from VEH/CON (\**p*<0.05, \*\**p*<0.01, \*\*\**p*<0.001); ^indicates significantly different from L-DE-71 (^*p* < 0.05). Bars and error bars represent values expressed as mean±s.e.m. n=5-8/group for epinephrine and *n*=8-21/group for BAT.

### DE-71 exposure alters hepatic glutamate dehydrogenase

Exposure to DE-71 increases blood glucose levels which may be due to enhanced hepatic glucose production. To examine this possibility, we tested the hypothesis that PBDEs increase the activity of glutamate dehydrogenase (GDH), a hepatic gluconeogenic enzyme. We found that exposure to L- and H-DE-71 significantly *reduced* enzymatic activity of GDH (One-way ANOVA: *Exposure effect* F_(2,33)_=0.8303, *p*=0.4448; Dunnett’s *post-hoc* VEH/CON vs L-DE-71 *p*=0.002, VEH/CON vs H-DE-71 *p*<0.0001, L-DE-71 vs H-DE-71 *p*=0.03) (**Fig. 7**). The effect of DE-71 was dose-dependent on this metric.

**Figure 7.**
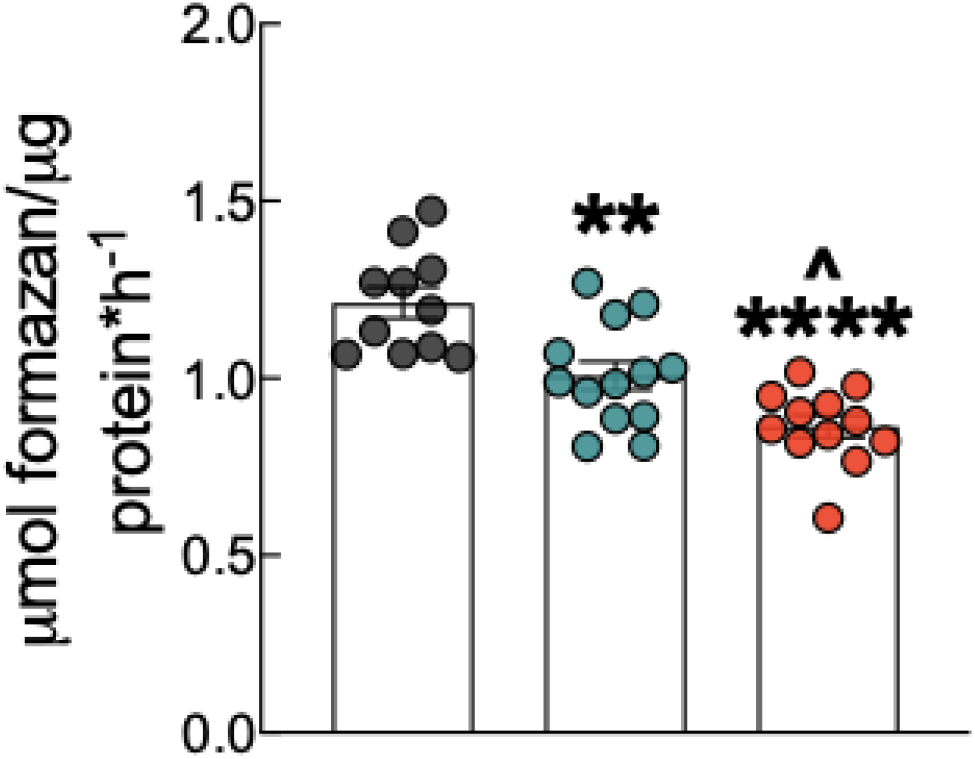
Effects of DE-71 on hepatic glutamate dehydrogenase (GDH). DE-71 exposure decreased GDH activity in a dose-dependent manner. *indicates significantly different from VEH/CON (\*\**p* < 0.01, \*\*\*\**p*<0.0001); ^indicates significantly different from L-DE-71 (^*p* < 0.05).Bars and error bars represent values expressed as mean±s.e.m. *n*=11-13/group.

### DE-71 exposure does not alter hepatic levels of endocannabinoids

Male F1 mice exposed to either dose of DE-71 displayed normal liver levels of the endocannabinoid (EC), anandamide (arachidonoylethanolamide, AEA), and related fatty acid-ethanolamides, docosahexaenoyl ethanolamide (DHEA), and n-oleoylethanolamide (OEA), a shorter analogue of AEA, when compared to VEH/CON (**Fig. 8A**). No changes were detected for the other primary EC, 2-arachidonoyl-sn-glycerol (2-AG), and related monoacylglycerol, 2-docosahexaenoyl-sn-glycerol (2-DG) (**Fig. 8B**).

**Figure 8.**
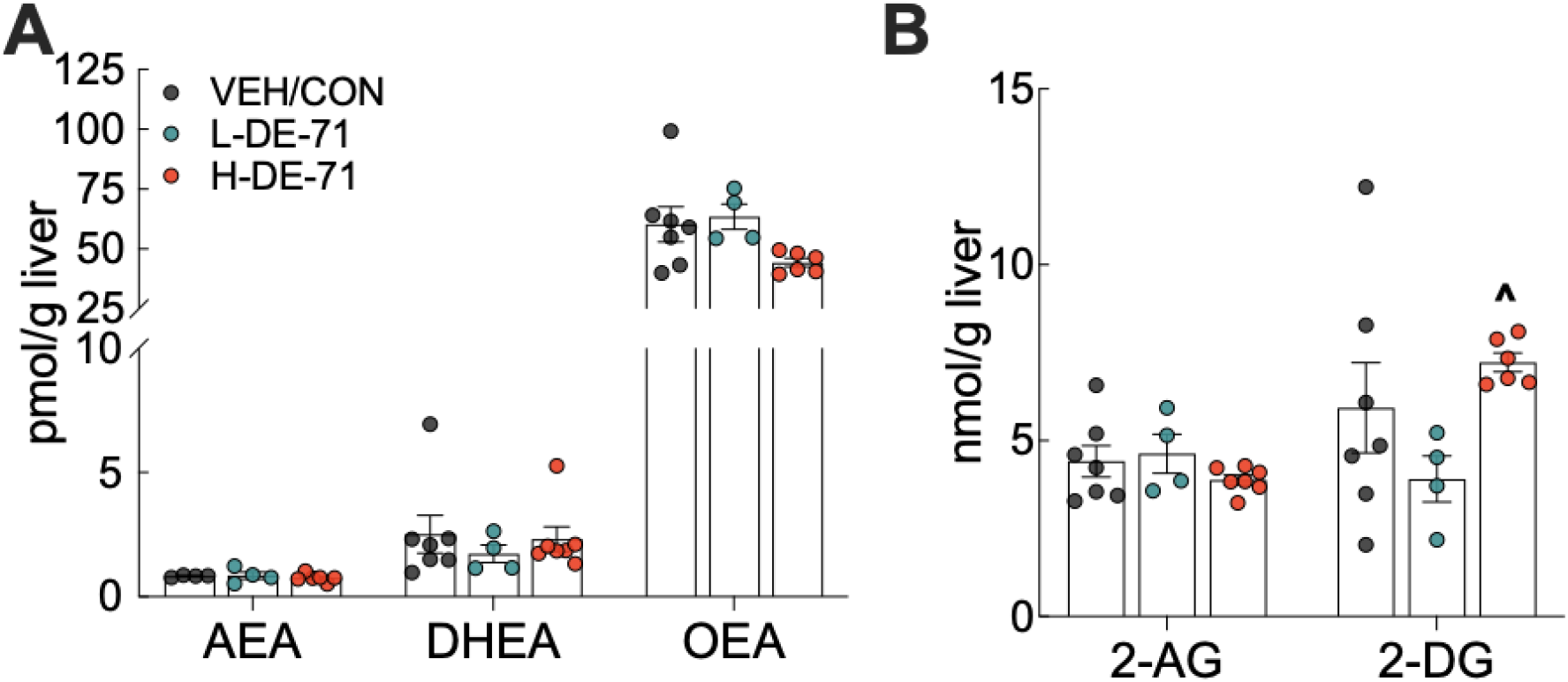
Hepatic levels of endocannabinoid (EC) and related fatty acid-ethanolamides in DE-71 exposed male mice. *Post mortem* liver tissue was analyzed using UPLC/MS/MS. (**A**) AEA, DHEA and OEA. (**B**) 2-AG and 2-DG. Bars and error bars represent values expressed as mean±s.e.m. *n*,4-7/group. AEA, arachidonoylethanolamide (Anandamide); DHEA, docosahexanoyl ethanolamide; OEA, n-oleoyl ethanolamide; 2-AG, 2-arachidonoyl-*sn*-glycerol; 2-DG, monoacylglycerol 2-docosahexaenoyl-*sn*-glycerol

## Discussion

Mounting evidence implicates environmental toxicants in increased susceptibility to obesity and diabetes (42), (43). Organohalogen compounds used inside the home, such as brominated flame retardants, are associated with metabolic reprogramming that may contribute to these and other diseases (44), (45). Previous findings from our lab demonstrate that female offspring exposed developmentally to a penta-mixture of PBDEs, DE-71, display a diabetogenic phenotype (23). In this article, we continue our examination of the relationship between early-life PBDEs and regulation of glucose homeostasis by examining male offspring. Here, we show that developmental DE-71 exposure produces altered glucose tolerance, glucose response after insulin challenge, glucoregulatory pancreatic and gut hormones and liver glucose production, and indices of sympathetic activity. Metrics studied were chosen based on *in vivo* and *ex vivo* biomarkers of diabetes in adulthood that are used to validate diabetic animals models (46). The reprogrammed metabolic profile observed in DE-71 exposed males does not exactly overlap with that reported for exposed female offspring (23).

### Human Relevance of the PBDE Dosing Paradigm

In our past works, we have demonstrated that exposure of mouse dams to 0.1 mg/kg/d and 0.4 mg/kg/d resulted in liver accumulation of 3.2 ppm in dam liver. This also resulted in 0.2-1.0 ppm in the liver and 0.07-0.3 ppm in brains of exposed female offspring (23), (24). The liver is of importance since it is a key organ for glucose homeostasis and lipid metabolism, while the brain levels may cause central disruption of glucose homeostasis (Kozlova et al., 2022, *In press*). Studies in toddlers report plasma ∑PBDE values with a range of 0.1-0.5 ppm (47), (48). Our liver levels in offspring are in range with maximum plasma levels reported in North American populations of Canadian indigenous Inuits and Crees (219–402 ng/g lipid wt) (20) and California U.S. women (~749.7 ng/g lipid wt) (49). In human studies, sum PBDE levels in children have been reported in the range of 43-366 ng/g lipid wt (48), (47), (50). Therefore, our dosing regimen produces a PBDE body burden that is similar to the levels experienced by certain human populations. Thus, exposures used in this study are relevant to human populations. In comparison, the EPA reference dose for penta mixture of PBDEs (DE-71) is 2×10^−3^ mg/kg/day or 2 ppb (37). However, reference dose is the minimum dose that does not produce bioactivity.

### Fasting Blood Glucose and Glucose Intolerance

In total, changes in glycemia metrics displayed by exposed male offspring represent the complex actions of PBDEs on the broad network of systems participating in the regulation of glucose metabolism total (51), (52). Extended fasting yielded reduced glycemia in DE-71 male offspring relative to VEH/CON possibly indicating dysregulated endocrine, hepatic and/or sympathetic contributions to glucose homeostasis (37). Results of the GTT conducted after 11 h fast duration showed that H-DE-71 exposure produces dramatic glucose intolerance as compared to VEH/CON. Interestingly, this phenotype was most prominent at H-DE-71 although L-DE-71 male offspring showed incomplete glucose clearance and/or utilization. These results suggest differential targets of DE-71 at 0.1 and 0.4 mg/kg. Despite the slight difference in dose of L- and H-DE-71, the chronic manner of exposure paradigm as well as the developmental timing may combine to augment responses in H-DE-71 to exogenous glucose challenge. Different doses of DE-71 within this tight range have been previously shown to produce differential bioactive actions (53). Indeed, in female offspring, we have previously shown that 0.1 mg/kg but not 0.4 mg/kg DE-71 produces glucose intolerance in GTT (23). In fact, a non-linear dose response has also been commonly reported on behavioral and cellular/molecular processes (24), (54).

### Altered Responses to Insulin Challenge

An ITT was used to examine insulin responsiveness in DE-71 exposed male offspring. In this experiments, developmental exposure to DE-71 at 0.1 mg/kg produced a significant reduction in recovery of plasma glucose clearance after 90-120 min post insulin i.p. challenge. In contrast, no significant differences in insulin K_ITT_ were detected and, instead, insulin sensitivity after initial challenge was normal as compared to VEH/CON. The combination of DE-71 effects on FBG, glucose tolerance and insulin responsiveness displayed by exposed male offspring demonstrates the complex actions of PBDEs on glucose homeostasis in males. Together, with DE-71 induced endocrine disrupting effects on glucoregulatory hormones (see below), maternal transfer of PBDEs in DE-71 during gestation and lactation produce metabolic reprogramming during development that may lead to metabolic disorders in adulthood. A developmental basis for adult disease has been proposed for metabolic and neurodevelopmental disorders (55). Indeed, body burdens of PBDEs are positively associated with T2D and MetS (21).

### Deregulated Pancreatic and Gut Glucoregulatory Hormones

In this study we showed that L-DE-71 males also display elevated glucoregulatory hormones GLP-1 and glucagon when compared to levels in VEH/CON. In contrast, insulin levels, which are sometimes altered in T2D (38), were similar across exposure groups. Glucagon secretion plays an essential role in the regulation of hepatic glucose production and may contribute to hyperglycemia typical of T2D. Gastrointestinal hormones like GLP-1, that are secreted in response to oral glucose, have incretin effects which were not seen here. Studies employing the GLP-1 receptor antagonist, exendin 9-39, indicate that endogenous GLP-1 performs important regulation over glucagon secretion during fasting and after a meal (56), (57). However, in the L-DE-71 exposed males, GLP-1 did not inhibit glucagon secretion since this group showed significantly elevated levels. The deregulation of GLP-1 and glucagon by PBDEs, may lead to a reduced antidiabetic effect of GLP-1, normally produced by inhibiting glucagon secretion, and may contribute to reduced glucose clearance seen in L-DE-71 exposed males. Other integrated effects of H-DE-71 may be responsible for glucose intolerance seen only at this dose. Nevertheless, plasma levels of these hormones may serve as an important marker of pre/diabetes in PBDE-exposed children. Indeed, clinical studies on T2D suggest that deregulation of GLP-1’s inhibitory effect on glucagon may be as important as GLP-1’s stimulatory effect on insulin secretion (39).

### Comparison to Glucoregulatory Phenotype in Exposed Female Offspring

The glucose metabolic profile in male offspring reprogrammed by DE-71 is not identical to that seen in exposed female offspring. Specifically, we detected sexually dimorphic differences in glucose homeostasis and insulin sensitivity. L-DE-71 exposed females showed glucose intolerance, insulin insensitivity and decreased insulin, decreased glucagon and no change to GLP-1(23). Therefore, perinatal DE-71 exposure in males resulted in different changes in these important components contributing to glucose metabolic regulation as compared to those characteristic of exposed female offspring. These differential profiles of DE-71 reinforce our previous findings showing sexual dimorphism in the regulation of lipid metabolism in exposed male and female offspring (Kozlova et al., 2022, *In press*). However, some actions of L-DE-71 (but not H-DE-71) were also observed in our report on exposed female, i.e., reduced hepatic glutamate dehydrogenase enzymatic activity, elevated adrenal epinephrine, and reduced thermogenic brown adipose tissue mass (23). Our findings should be leveraged to help inform risk management analysis, provide best treatments in the clinic and improve consistency in studies on experimental animals. The long-lasting changes in glucose and lipid metabolism may be a risk factor for developing T2D and liver steatosis with increasing prevalence in the US population (58).

### DE-71 alters hepatic enzymatic and and sympathetic markers of activity

The sympathetic nervous system exerts regulatory control over nutrient release and utilization. We measured brown adipose tissue mass and adrenal epinephrine as indices of sympathetic activity. Brown adipose tissue (BAT) activity (41) increases energy expenditure and has been inversely associated with diabetes and fasting glucose level (41). Here we report reduced BAT mass in L-DE-71 relative to VEH/CON, which may have implications for glucose homeostasis. Adrenal epinephrine, which is regulated by and responds to sympathetic nervous system activation, was elevated in L- and H-DE-71, a profile that is characteristic of MetS (59). Glutamate dehydrogenase activity measured in exposed male livers was reduced in L- and H-DE-71. Interestingly, DE-71 exposure produced similar results on BAT, adrenal epinephrine and hepatic glutamate dehydrogenase enzymatic activity in exposed female offspring (Kozlova et al., 2022, *In press*).

## Conclusion

Altogether, the current results determined from *in vivo* and *ex vivo* parameters in DE-71-exposed male offspring indicate broad, complex effects of developmental exposure to PBDEs on glucose homeostasis and glucoregulatory parameters involving various systems. Importantly, our findings warn that maternal transfer of PBDEs can produce metabolic and endocrine reprogramming of male offspring that persists into adulthood and may contribute to metabolic diseases. Both exposed male and female offspring are susceptible, albeit they display different and complex phenotypes. We hope that our findings will inform risk management determination, clinical practices and study design in animal studies to uncover further the participation of the environment.

## Declarations

### Funding

This work was supported by UC MEXUS small grant (M.C.C.), a University of California UC-Hispanic Serving Institutions Doctoral Diversity Initiative (UC-HSI DDI), President’s Pre-Professoriate Fellowship and Dept. of Education (UCR STEM-HSI) award (E.V.K.); NIH grants DK119498 and DK114978, and TRDRP grant T29KT0232 (N.V.D.).

### Conflicts of interests/Competing interests

The authors declare that the research was conducted in the absence of any commercial or financial relationships that could be construed as a potential conflict of interest.

### Ethics approval

Care and treatment of animals was performed in accordance with guidelines from and approved by the University of California, Riverside Institutional Animal Care and Use Committee (IACUC) (AUP# 20170026 and 20200018).

### Statement on studies involving human subjects

No human studies are presented in this manuscript.

### Inclusion of identifiable human data

No potentially identifiable human images or data is presented in this study.

### Data availability statement

The raw data supporting the conclusions of this article will be made available by the authors, without undue reservation.

### Consent for publication

All authors reviewed and approved the final manuscript.

### Disclaimer

The opinions and assertions expressed herein are those of the author(s) and do not necessarily reflect the official policy or position of the Uniformed Services University or the Department of Defense.

J.K. is now a 2nd Lieutenant at the Uniformed Services University, Department of Defense. Her work was performed at the University of California, Riverside before becoming a military officer. However, we want to emphasize that the opinions and assertions expressed herein are those of the authors and do not necessarily reflect the official policy or position of the Uniformed Services University or the Department of Defense.

### Author Contributions

Conceptualization, M.C.C., E.V.K.; Methodology, M.C.C., E.V.K., N.V.D., J.M.K., B.D.C.; Validation, M.C.C., E.V.K., B.D.C., P.A.P., N.V.D.; Formal Analysis, E.V.K., M.C.C., P.A.P., N.V.D.; Investigation, E.V.K., B.D.C., P.A.P., J.M.K., A.E.B., M.C.C, M.E.D.; Writing – Original Draft, E.V.K., M.C.C., A.E.B.; Writing – Review & Editing, M.C.C., E.V.K., A.E.B; Visualization, E.V.K.; Resources, M.C.C., N.V.D.; Data Curation, E.V.K., M.C.C.; Supervision, M.C.C., E.V.K. N.V.D. Project Administration, M.C.C., E.V.K., Funding Acquisition, M.C.C., E.V.K., N.V.D. All authors reviewed and approved the final manuscript.

## Abbreviations

2-AG: 2-arachidonoyl-sn-glycerol
2-DG: 2-docosahexaenoyl-sn-glycerol
AEA: arachidonoylethanolamide
AUC: area under curve
BAT: brown adipose tissue
BDE(s): decabromodiphenyl ethers
DE-71: pentabrominated diphenyl ether mixture
DHEA: docosahexaenoyl ethanolamide
EC: endocannabinoid
EDC(s): endocrine disrupting chemicals
EDTA: ethylenediamine tetraacetic acid
EIA/ELISA: enzyme-linked immunosorbent assay
EPA: Environmental Protection Agency
FBG: fasting blood glucose
GDH: glutamate dehydrogenase
GLP-1: glucagon-like peptide 1
GTT: glucose tolerance test
ITT: insulin tolerance test
LOEL: lowest-observed-adverse-effect level
MDC(s): metabolism-disrupting chemicals
MetS: metabolic syndrome
OEA: n-oleoylethanolamide
PBDE(s): polybrominated diphenyl ethers
PND: postnatal day
POPs: persistent organic pollutants
T2D: type 2 diabetes
UPLC/MS/MS: ultra-high performance liquid chromatography with tandem mass spectrometry
VEH/CON: vehicle control group

## Acknowledgements

We are grateful to ALPCO for the gift of GLP-1 and Glucagon immunoassay kits. We also thank Dr. M. Valdez, Dr. G. Gonzalez and Karthik Basappa (Curras-Collazo Lab), Dr. P. Deol and J. Evans (Sladek Lab) and Dr. Donovan Argueta and Andrea Dillon (DiPatrizio lab) for their technical assistance. We are grateful to Great Lakes Corporation for the gift of DE-71 and to Drs. F. Sladek and I. Ethell for the gift of C57BL6 mice. We thank J. Phan for assistance with animal husbandry. We thank Dr. Sargeet Gill for the use of plate reader.

## Notes

### Competing Interest Statement

The authors have declared no competing interest.

